# Generating realistic artificial Human genomes using adversarial autoencoders

**DOI:** 10.1101/2023.12.08.570767

**Authors:** Callum Burnard, Alban Mancheron, William Ritchie

## Abstract

A publicly available human genome serves as both a valuable resource for researchers and a potential risk to the individual who provided the genome. Many actors with selfish intentions could exploit it to extract information about the donor’s health or that of their relatives. Recent efforts have employed artificial intelligence models to simulate genomic data, aiming to create synthetic datasets with scientific merit while preserving patient anonymity. However, a major challenge arises in dealing with the vast amount of data that constitutes a complete human genome and the resources required to process it.

We have developed a dimension reduction method that combines artificial intelligence with our knowledge of in vivo mutation association mechanisms. This approach enables the processing of large amounts of data without significant computational resources. Our genome segmentation follows chromosomal recombination hotspots, closely resembling mutation transmission mechanisms. Training data is sourced from the 1000 Genomes Project, which catalogues over 2500 genomes from diverse ethnic groups. Variational autoencoders, utilising neural networks, serve as an extension to the generative model. Wasserstein Generative Adversarial Networks (WGAN) are a benchmark among generation methods for various data types.

After optimisation of our data simulation strategy our pipeline allows the generation of a simulated population meeting several essential criteria. It demonstrates good diversity, closely resembling that found in the reference dataset. It is plausible, as newly generated combinations of mutations do not disrupt the linkage disequilibria found in humans. It also preserves donor anonymity by synthesising combinations of reference genomes that are distant from reference samples.

## Introduction

The sequence of nucleotides in the DNA contained in our cells constitutes the biological information necessary to characterise us as individuals. This information covers aspects as innocuous as the colour of our hair, eyes and skin, as well as functional elements of our bodies like our capability to digest certain foods, and can go so far as to cause diseases. The progresses in sequencing technologies over the past two decades allow us to express these characteristics as variations from a reference genome, established by studying large swathes of the population (Schneider et al. 2017).

We are all uniquely identified by these aspects, and it is becoming increasingly easy for healthcare services, research teams and private companies to identify the genomic mutations underlying the phenotypes we observe. This leads to a tradeoff, where this data can be used to better treat diseases, in particular when dealing with personalised medicine, but at the same time could have negative repercussions on the everyday lives of the bearers of these variations (Joly, Ngueng Feze, and Simard 2013; Prince 2017; Price and Cohen 2019). Great care is therefore needed when dealing with this data, in particular when sharing it with other actors.

A large number of regulations are active across the world regarding the storage and sharing of medical data, especially in the European Union, where the authors are based (Shabani and Marelli 2019). However, it is much more complex to protect this data when it is in use. Methods such as homomorphic encryption (Kim and Lauter 2015) or federated learning (Kuo and Pham 2021) could potentially alleviate privacy issues, but they are either very costly to implement or reduce the information that can be extracted from sequencing data.

A different approach to the privacy issue is to synthesise novel genomic data not directly derived from an individual’s sample. In the past, these methods have often focused on the simulation of an entire population and the coalescence of alleles within it (Balloux 2001; Tahmasbi and Keller 2016; Carvajal-Rodríguez 2008; Peng and Kimmel 2005). In recent years, teams have sought to use available data describing the frequency of mutations in populations and create novel samples by assigning these mutations in novel combinations (Juan et al. 2020; Yue and Liti 2019; Mu et al. 2015). One of the most recent projects of this type uses artificial intelligence methods, specifically generative approaches to determine appropriate combinations of mutations for sample generation (Yelmen et al. 2021). This approach produces diverse, realistic and novel populations of genomes.

The main limitation however of contemporary AI methods is the size of both the training data and generated genomes. A classical artificial neural network is densely connected, meaning each neuron in a given layer is connected to each neuron of the following and previous layers. Increasing the size of the sample used for training the model therefore leads to exponential increases in calculation cost.

### Graphical abstract

We present here the Haplotypic Human Genome Generator (H2G2), a method to generate human genomic data on an increased scale. We first segment the data based on available biological knowledge and create a compressed representation of these subsections of data using dimension reduction methods. Then, these genomic identities are processed by a generative neural network to simulate novel samples, while remaining coherent with the source dataset.

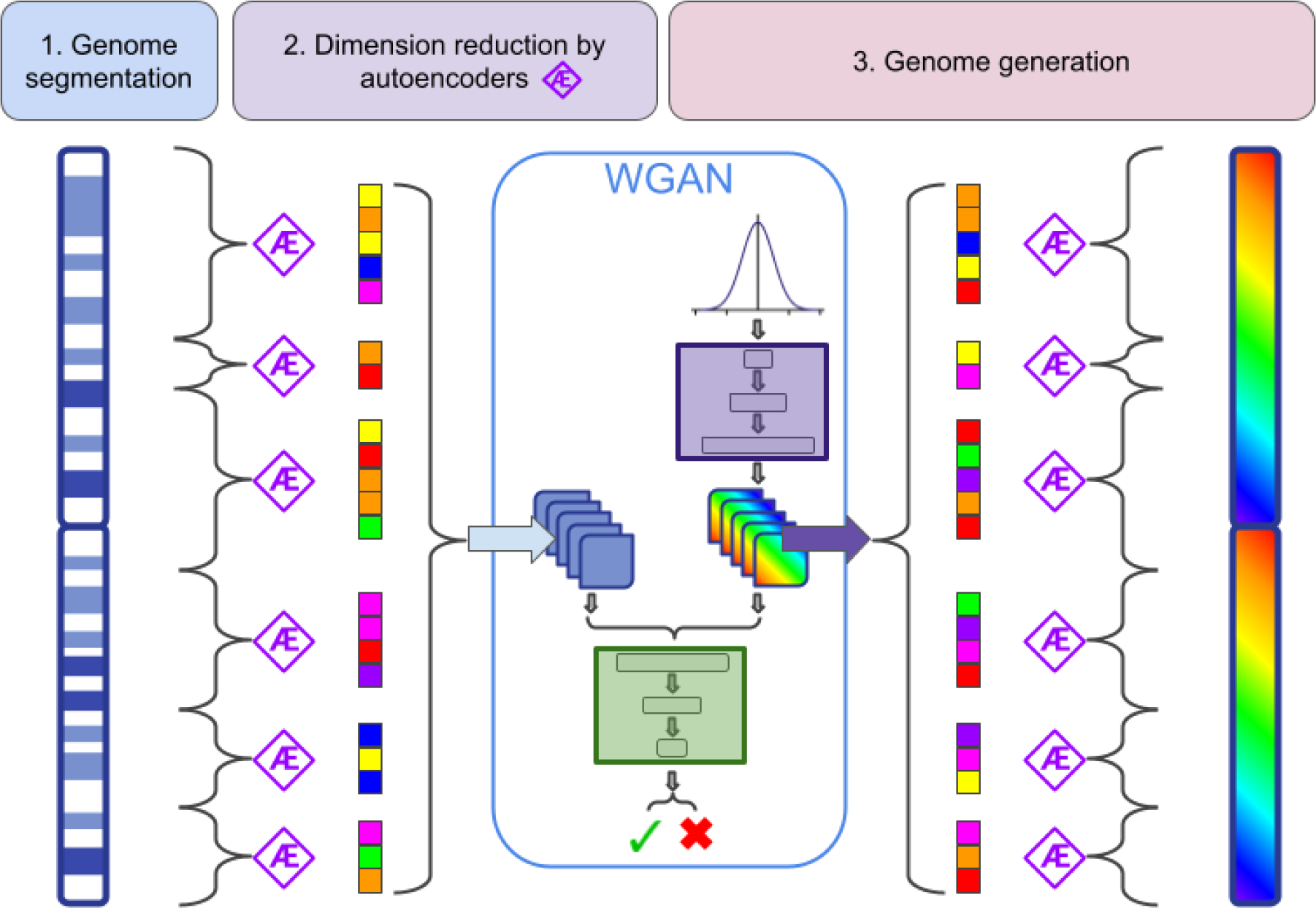
Pipeline for increased scale of synthetic genome generation. Genomic data is segmented into subsections, then compressed using deep autoencoders. Finally a generative neural network learns from this compressed genomic data to create novel encoded samples, which are then decoded back into full samples.

We use Human genomic data from the 1000 Genomes Project (The 1000 Genomes Project Consortium 2015), realigned onto HG38 (Lowy-Gallego et al. 2019), for its availability, high global diversity and standardised methodologies. Mutation data for chromosome 1 was segmented based on hotspots of recombination likelihood as determined by a high resolution map of recombination events established across thousands of detailed samples (Halldorsson et al. 2019). These recombination events dictate which mutations tend to be inherited together across generations, and therefore orchestrate the biological linkage between genomic variations at a large scale. Following this logic, mutations within a subsection share higher correlation than mutations contained in two different subsections, and allow for a more accurate compression of the data by dimension reduction methods. We chose autoencoders to compress the data. These are a family of unsupervised machine learning methods using artificial neural networks to create latent representations of samples by compressing them more and more at each layer in the network (Kramer 1992; Kingma and Welling 2013). The models also implement a decoder to restore the compressed data into its full state, as faithful to the original samples as possible over the course of the training process. Their strengths in dealing with any type of data, as well as handling nonlinear datasets with variable multimodal distributions, are particularly relevant for this project.

The generation method we chose was the Wasserstein GANs (Arjovsky, Chintala, and Bottou 2017). Their robust training process, by avoiding mode collapse and being less sensitive to hyperparameters when compared to other similar models, is particularly relevant for a project that seeks to increase the scale of data to be simulated.

Additionally, we validated the realism of our simulated population by applying Chromopainter to reference and synthetic genomes (Lawson et al. 2012). Meta-characteristics of these two groups were calculated based on its output to compare them and highlight their similarities.

## Results

### Genome segmentation

The first step of our approach is to segment the genome into portions that are amenable to compression using autoencoders. To this end we compared two segmentation approaches which were a linear segmentation of the genome into equal sized bins and a segmentation based on crossing-over hotspot regions. Recombination hotspots were used to segment mutation data for HG38 chromosome 1 (supp. figure 1a), setting minimum and maximum values for number of mutations per subsection (supp. figure 1b). This method was compared to using a naive method of simply separating mutations into bins of equal size. Weight pruning and latent space reduction were also applied to these models to maximise encoding efficiency (supp. figure 1c)

**Figure 1:**
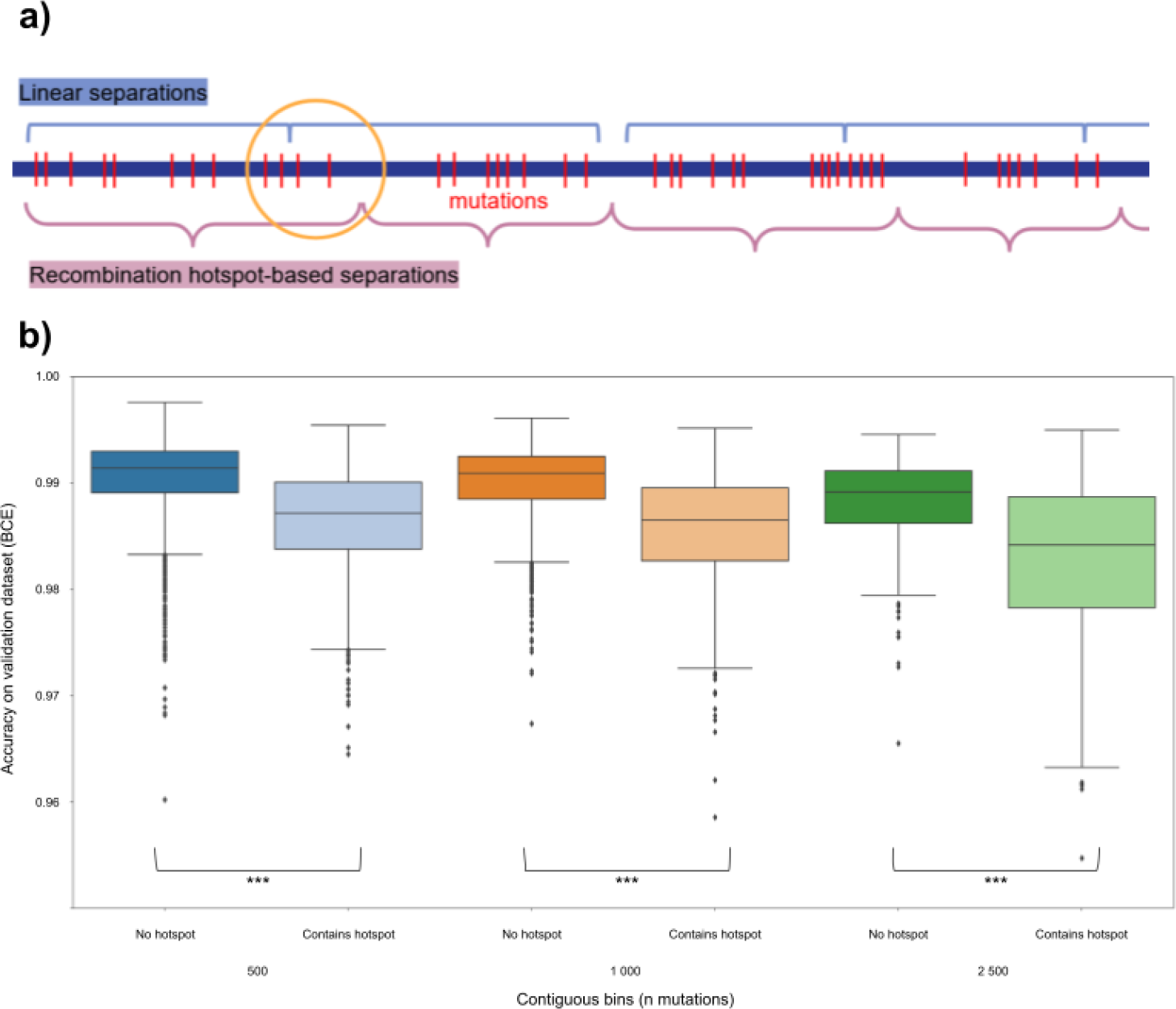
Genome segmentation. Recombination hotspots provide higher accuracy segmentation. a)Our model explaining linear separations splitting up biological links between mutations. Mutations that share more biological meaning with their neighbours on one side than on the other are sometimes attributed to the wrong group of mutations when not using recombination hotspot based methods, leading to less accurate reconstruction by AE. b)Boxplot showing the accuracy of autoencoder models applied to contiguous bins of mutations. For each size of bins, they were separated into bins containing a high intensity hotspot (light colour) and those that don’t (dark colour). Statistical significance of the differences were tested using Mann-Whitney U’s test.

According to our model (figure 1a), some mutations can become separated from others that share the most biological significance with them by chance when using a naive approach, whereas the recombination hotspots-based method preserves these links. Analysing bins of mutations containing a strong hotspot (figure 1b) confirms this, as these sections that are split by a recombination hotspot show significantly lower accuracy after autoencoding when compared to sections that do not contain a strong recombination hotspot.

We selected using hotspot-based segregation of mutations for the rest of this study, allowing for a minimum of 500 and a maximum of 5000 mutations per subsection.

### Dimension reduction by autoencoders

Segments generated from the previous section were then compressed using autoencoders (see methods section for full details). We first investigated the frequency of each mutation before and after compression via the autoencoders. We discovered that mutations with a low frequency in the reference dataset tend to be missing from the decoded dataset (figure 2a, upper left plot, highlighted by red box). This can be due to the tradeoff between higher compression rates and encoding rare variations. The autoencoder is not learning the rare occurrences in which the mutation should be present as a reasonable tradeoff for a gain in compression The poor representation of rare mutations could also be due to neurons in the model “dying”, which can happen when using the ReLU activation function (Ramachandran, Zoph, and Le 2017). We therefore investigated multiple alternative activation functions, as well as a Variational AutoEncoder (VAE) (Kingma and Welling 2013). Results in figure 2a show that different activation functions do indeed affect the number of disappearing mutations. Interestingly, the TanH activation function yields the fewest of these disappearing mutations, but also systematically tends to overrepresent the frequency of mutations in the dataset (figure 2a, bottom row, highlighted in red).

**Figure 2:**
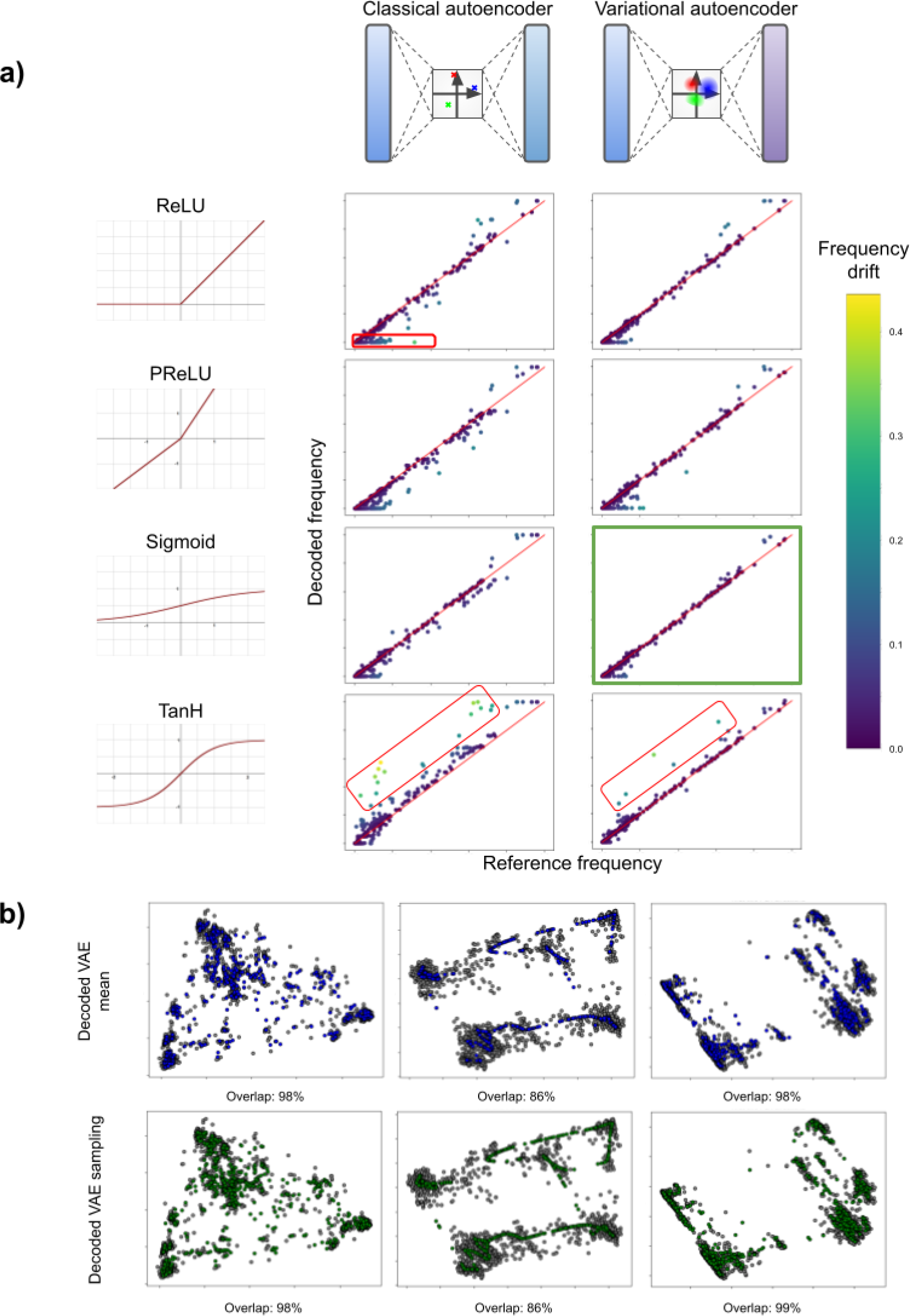
Dimension reduction by autoencoders. Model and hyperparameter optimisation for accurate reconstruction. a)Scatter plots comparing mutation frequencies in the reference dataset to the frequency of that mutation in the dataset after encoding and decoding by various autoencoder models. Occurrences where frequencies are equal align with red diagonal. Classical autoencoders and variational autoencoders were tested, each with four different activation functions. For each plot, the colour of the dot indicated the frequency drift, which is simply the absolute difference of the two frequencies. Particular drift patterns observed across different setups are highlighted manually. b) Scatter plots of Principal Component Analyses (PCA) of various datasets of genomic subsections. Each column is a different genomic subsection. In each plot, the reference dataset is shown as grey dots, and a dataset obtained after encoding then decoding by a Variational AutoEncoder (VAE) is shown in colour. Top row shows data obtained by decoding the mean of the projection of each individual in its latent space. Bottom row shows data obtained from the same VAE, but using a sampling around the mean of the projection of each individual in its latent space.

Overall, using a VAE with either sigmoid or TanH activation function show the lowest number of disappearing mutations as well as a low frequency drift (supp. figure 2a). Sigmoid activation function was chosen, so as to avoid the visible bias of highly overrepresented mutations that occur when using TanH, both in VAE and classical AE.

PCA projections show that these VAE models are very good at reconstructing the population after dimension reduction (figure 2b and supp. figure 2b). Some do exhibit mode search, where the model simplifies the information it learns down to a set number of modes which, once decoded, map to the main modes of the reference distribution.

### Genome generation

Following the segmentation and compression steps described above, we trained a Generative Adversarial Network using Wasserstein loss (WGAN) on encoded subsections of genomic data spanning over 15000 mutations, equivalent to 1 megabase of DNA. It produces simulations of encoded genomic data. After decoding this data and applying the same PCA projection, we can see that the samples generated by this model remain realistic, as their coverage of the reference data is similar to that of the decoded data in figure 2b, and explore some space which the VAE did not utilise completely (figure 3a and supp. figure 3).

**Figure 3:**
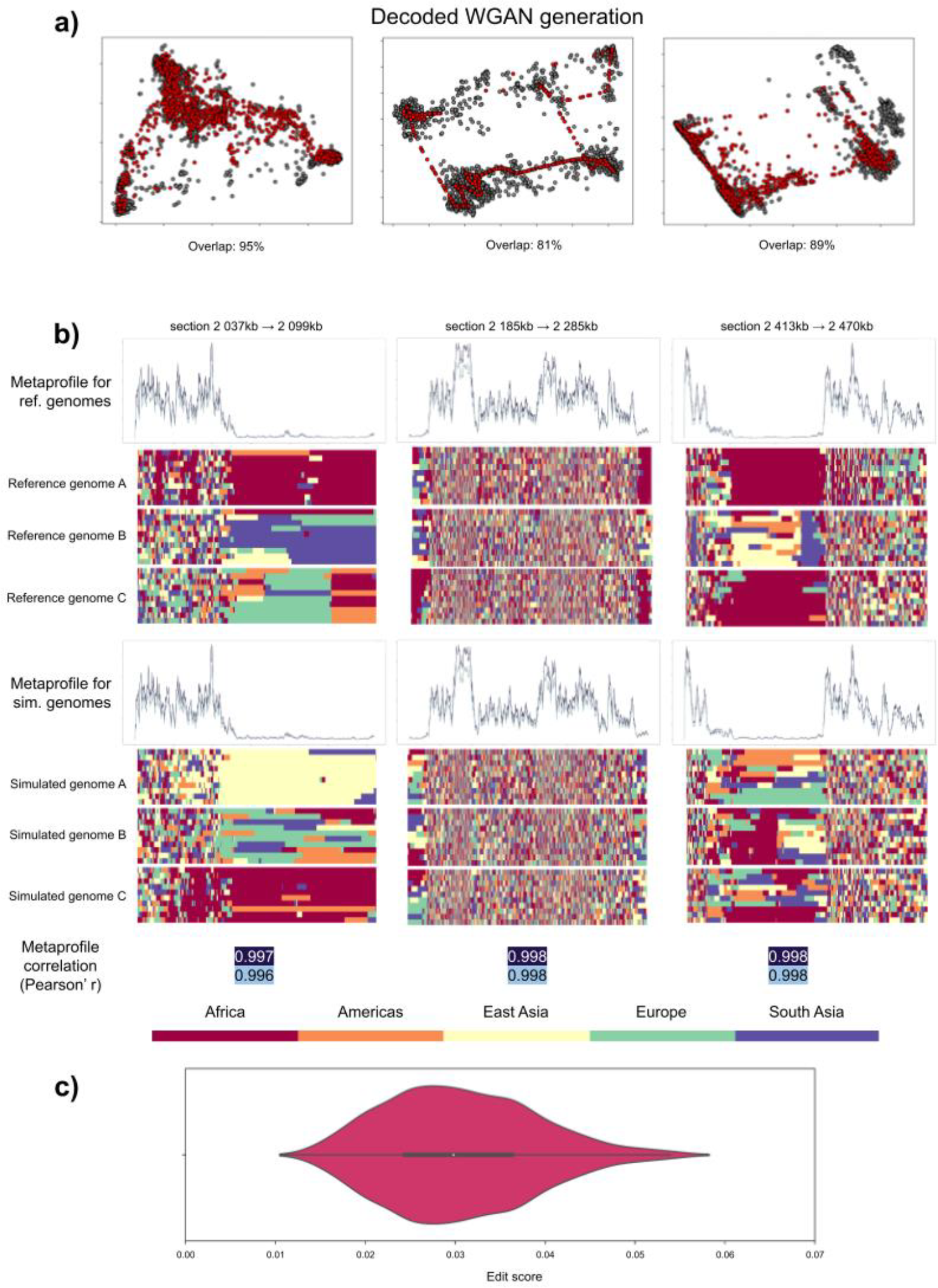
Genome generation. Realistic and novel genomes are simulated using a multi-step pipeline. a)Scatter plots of Principal Component Analyses (PCA) of data generated by Generative Adversarial Network using Wasserstein loss (WGAN) in red, versus reference data in grey for three genomic subsections. b)Chromopainter metaprofiles calculated over all samples for three genomic subsections, and three corresponding Chromopainter illustrated reconstructions from the datasets used to construct the metaprofiles. Chromopainter gives as output a series of reconstructions for each sample indicating the most probable ancestry for each mutation in the sample. Each plot is a different sample, each row in each plot is a reconstruction by Chromopainter. The horizontal axis is each mutation in the genomic subsection. Metaprofiles show how frequently this ancestry changes within samples. In dark blue, ancestry change from one individual to another, in light grey, ancestry change from one population to another. c)Boxplot of edit score (based on Hamming distance) for each generated sample against the reference dataset, measured across all genomic subsections combined.

To analyse the validity of our simulated genomes we used a population based approach where we estimate how well the ancestry composition of the simulations match that of the real genomes. To this end we used a tool called Chromopainter which reconstitutes the genomic ancestry of a query population based on the mutation profile of a donor population. Chromopainter was applied to both 78 reference genomes (three per sub-population listed in (Lowy-Gallego et al. 2019)) and 100 simulated genomes. From the raw results obtained, we established metaprofiles of ancestry switches, based on all reference or simulated genomes analysed (figure 3b). These metaprofiles show strong correlation between the reference and simulated datasets. This means that our simulated population could realistically be obtained through population admixture *in vivo*.

One other characteristic to analyse for this new population is privacy loss - that is, how close are these simulated genomes to the ones found in the reference dataset? To this end we use the Hamming distance to measure the number of substitutions necessary to transform one sequence into another of same length (Hamming 1950). It was used here as a part of the edit score, which is the ratio between the Hamming distance from a simulated genome to its nearest neighbour in the reference dataset and the total length of the sequence. This score therefore ranges from 0 (identical sequences) to 1 (no common elements in sequence). Applying it to the simulated section of chromosome 1 (figure 3c) shows that none of our simulated genomes are exact copies of any reference samples. The minimum score is 0.01, which is equivalent to just over 150 mutations difference.

## Discussion

In this study we propose a novel approach for generating human genomic data at a large scale and to a high degree of detail. This pipeline combines a novel dimension reduction approach, compression and a GAN structure. These allow a high level of parallelisation and a reduction of computational resources required to generate genomes. Specifically, we show that our models can generate realistic and novel mutation data at a larger scale and at an increased level of detail than previous AI based methods (Yelmen et al. 2021). One of the main applications we envision for this sample generation pipeline is to create synthetic cohorts for specific sets of patients. A WGAN model that has been trained to process general genomes could be specialised on a smaller dataset, obtained for example by a healthcare establishment sequencing all its patients bearing a disease, or exhibiting similar traits. This generator would then create novel samples that follow the same implicit rules as the original patients in the cohort. This would allow the sharing of the cohort’s general information, which could be an important resource for research teams, without revealing the identity of any one person in it.

There are many other potential use cases that one could imagine for realistic synthetic genomes generated by AI. Creating a digital twin of an individual or a population could be a useful asset for sharing or experimenting on. Augmenting already available datasets of genomes might lend more robustness to certain research projects. Finally, the autoencoding of genomic data and its accurate reconstruction is a potential method of processing these samples in a secure manner, through federated learning or homomorphic encryption.

However, combining sequencing data that was obtained using different techniques will require caution as to the standardisation methods used. Additionally, we have limited ourselves to short variations, whereas many large scale, structural, and/or copy number variations also have large impacts on the phenotype of the organisms bearing them.

The haplotypes used here as samples are based on available phasing data, which is not guaranteed to be completely accurate, especially at long range. It would be beneficial to projects like these to resolve long range links between mutations, so as to present high accuracy samples to our ML models. Haplotypes are also not consistent between chromosomes, as it is complex to establish biological linkage between mutations across chromosomes, but future applications may look to tie data from different chromosomes together. A project that is able to provide correlation data across chromosomes for certain populations would enable this scale to be explored.

The recombination loci used to segment the genome here is based off of a study focused on an icelandic population. This represents a very small subset of the Human population, and therefore the hotspots identified are not accurate on a global scale. They still constitute a significant gain of reconstruction accuracy for this methodology and a good starting point, but a similar study on a larger scale would be very beneficial in establishing more widespread recombination hotspots.

Regarding the machine learning models implemented in this project, a great deal of exploration remains possible, and our methods are by no means set in stone. VAEs using sigmoid activation function were shown to be the best dimension reduction model amongst those we tested, but there are many more architectures that could be tested in search of even greater efficiency and accuracy. As for generative models, these past years have shown remarkable progress in stable diffusion-based models that allow for detailed illustrations based on user specifications. Although these models are based on convolution neural networks, which are particularly well suited for processing images, which we do not use, it would be interesting to study their application to genomic datasets.

Evaluating the realism of a synthetic genome is a difficult task, in part because no standardised tools or methods exist for this purpose, and even more so for a synthetic population. The criteria proposed here - PCA tiling overlap score, Chromopainter metaprofile correlation and edit score all highlight various aspects of a simulated population that are desirable. To go further, it could be interesting to investigate the biological relevance of mutations contained in these simulated genomes and seek to measure degrees of similarity and difference using variations with known biological meaning.

## Methods

### Python environment

Unless specified otherwise, all scripts were run in python 3.8.10, using numpy 1.24.3, pandas 1.2.4 and keras 2.12.0 with tensorflow backend. Scripts run on the Genotoul cluster were launched using SLURM 22 and parallel release 20180122 (Tange 2011).

## Code availability

The main scripts used during this project for data preprocessing, building ML models and the custom training loop for the WGAN model, are available at this github repository: https://github.com/callum-b/H2G2/

### Data

Mutation data for 2504 individuals as well as donor information comes from the 1000 Genomes Project (phase 3), realigned onto HG38 in (Lowy-Gallego et al. 2019). It was downloaded here: http://ftp.1000genomes.ebi.ac.uk/vol1/ftp/data_collections/1000_genomes_project/release/20190312_biallelic_SNV_and_INDEL/

Each donor genome was separated into haplotypes. Mutations on chromosome 1 were filtered to only include those present in at least three of these haplotypes, so as to remove data with little detectable correlation with the rest of the dataset. Contiguous bins of a set number of mutations were then created from this list of variants.

Crossing over data was obtained from a high definition map of recombination rates throughout the human genome. Ranges for chromosome 1 were extracted, and those with a recombination rate of <=5 cM per Mb were removed to eliminate noise. The remaining ranges were reduced to their centres and sorted by recombination rate. Any low-intensity ranges within 5kb of a higher intensity range were removed. This resulted in approximately 2500 hotspots across all of chromosome 1.

Splitting mutation data into sections bounded by hotspots use these delimiters. Starting at one end of chromosome 1, mutations were added to a section until a hotspot is encountered, at which point the data collected thus far is saved to a file and a new section begins. If fewer than a minimum threshold of mutations are encountered, instead these mutations are appended to the previous section. If more than a maximum threshold of mutations are encountered, an “artificial hotspot” is created, and data is saved to disk for this range.

### Autoencoders

Classical autoencoders are composed of an encoder with three hidden layers, then a bottleneck, and finally a mirrored encoder setup for the decoder. Each layer uses ReLU (unless otherwise specified) for output, and there is a 20% dropout rate in between the hidden layers in both encoder and decoder. Bottleneck size is defined as input size divided by 100, rounded up, and the size of the hidden layers is calculated using geometric spacing. It uses classical reconstruction loss, measuring binary cross entropy between the input data and the reconstruction data.

Variational autoencoders are composed of an encoder with three hidden layers, then a bottleneck and finally a mirrored encoder setup for the decoder. Each layer is connected by sigmoid (for the final models used for compressing data), except for the bottleneck, which uses linear activation, and the output layer, which uses ReLU with a maximum value of 1. There is a 40% dropout rate in between each hidden layer, for the encoder and the decoder. Bottleneck size is defined as input size divided by 100, rounded up, and the size of the hidden layers is calculated using geometric spacing. Its loss is measured as the sum of classical reconstruction loss and the Kullback-Leibler divergence (Kullback and Leibler 1951), to ensure a locally continuous latent space.

The training process for both these model architectures uses a training set and a validation set, with an 80/20 split. The models are trained while monitoring loss on the validation dataset, and when this value increases (with a patience of 10 epochs, or 30 when using sigmoid and tanH activation due to slower convergence), training is halted and the weights of the model are restored to its best performing iteration. The accuracy of this saved model on the validation dataset is used as its performance metric in the figures presented here.

## WGAN

The Generative Adversarial Network used here is composed of two networks: a critic, that assigns a realness score to each sample it analyses, and a generator, that creates novel samples that attempt to fool the critic. It implements the Wasserstein (earth mover) loss function, as well as a custom training loop so as to train the critic more than the generator, as per the usage of Wasserstien loss recommends. This loop also allows checkpoints to be saved during training.

The latent vector used contains 1000 random variables drawn from a Gaussian distribution with mean 0 and variance 1. The generator is composed of three hidden layers, using ReLU activation, their sizes determined by the total number of latent dimensions to predict, depending on the genomic sections processed. The first hidden layer is 10 times smaller, the second hidden layer is 5 times smaller, and the third hidden layer is 2 times smaller than that total. The output layer comes after this, using the same size and activation function as the combined latent spaces of the autoencoders.

The critic is also composed of three hidden layers and an output, using ReLU activation, and their sizes also depend on the total number of latent dimensions considered. The first hidden layer is 10 times smaller, the second is 50 times smaller, and the third is 100 times smaller. The output layer is a single neuron with linear output.

The models, both the WGAN as a whole and the critic individually, are trained using RMSProp optimiser with a learning rate of 0.00005. In each training loop, the critic is trained 5 times more than the generator. We are using Wasserstein distance, so the aim of the critic is to maximise the distance between the real samples and the simulated ones, and the aim of the generator is to minimise this distance. To implement this during training, the loss of the critic is its loss on simulated samples minus its loss on real samples, and the loss of the generator is equal to -1 times the loss calculated by the critic on the samples created by the generator for this training loop.

Multiple checkpoints of the model were sampled during training, and the data they produced were evaluated to find the optimal stopping point. Checkpoints were saved every 10,000 training epochs, and the results presented here are taken from the model trained for 30,000 epochs.

### Overlap score

Overlap is calculated based on a tiling of the PCA projection space. For each sample in one dataset, if its tile also contains a sample from the other dataset, it is considered to overlap. This process is applied to both datasets then averaged.

### Chromopainter

We estimated the ancestral diversity of our simulated genomes using Chromopainter, which attempts to assign sections of donor genomes to query genomes. We use this as a verisimilitude measure, as our simulated genomes should present approximately the same profile of crossing overs as the reference genomes.

We applied this tool to a reference query dataset, composed of three haplogenomes from each subpopulation listed in the 1000 Genomes Project phase 3 supplementary information, and to 100 randomly chosen simulated haplogenomes, for three genomic subsections. The donor dataset is composed of the 1000 Genomes Project individuals, without the members of the reference query dataset.

### Edit score

The edit score used here to compare simulated populations to reference populations is based on the Hamming distance metric. We measure this distance for each simulated individual to each reference individual and retain the lowest of these values, which is then divided by the length of the sequence. This creates an “edit rate”, which allows comparison between sections of different sizes.

## Supplemental figures

**Supplementary fig 1:**
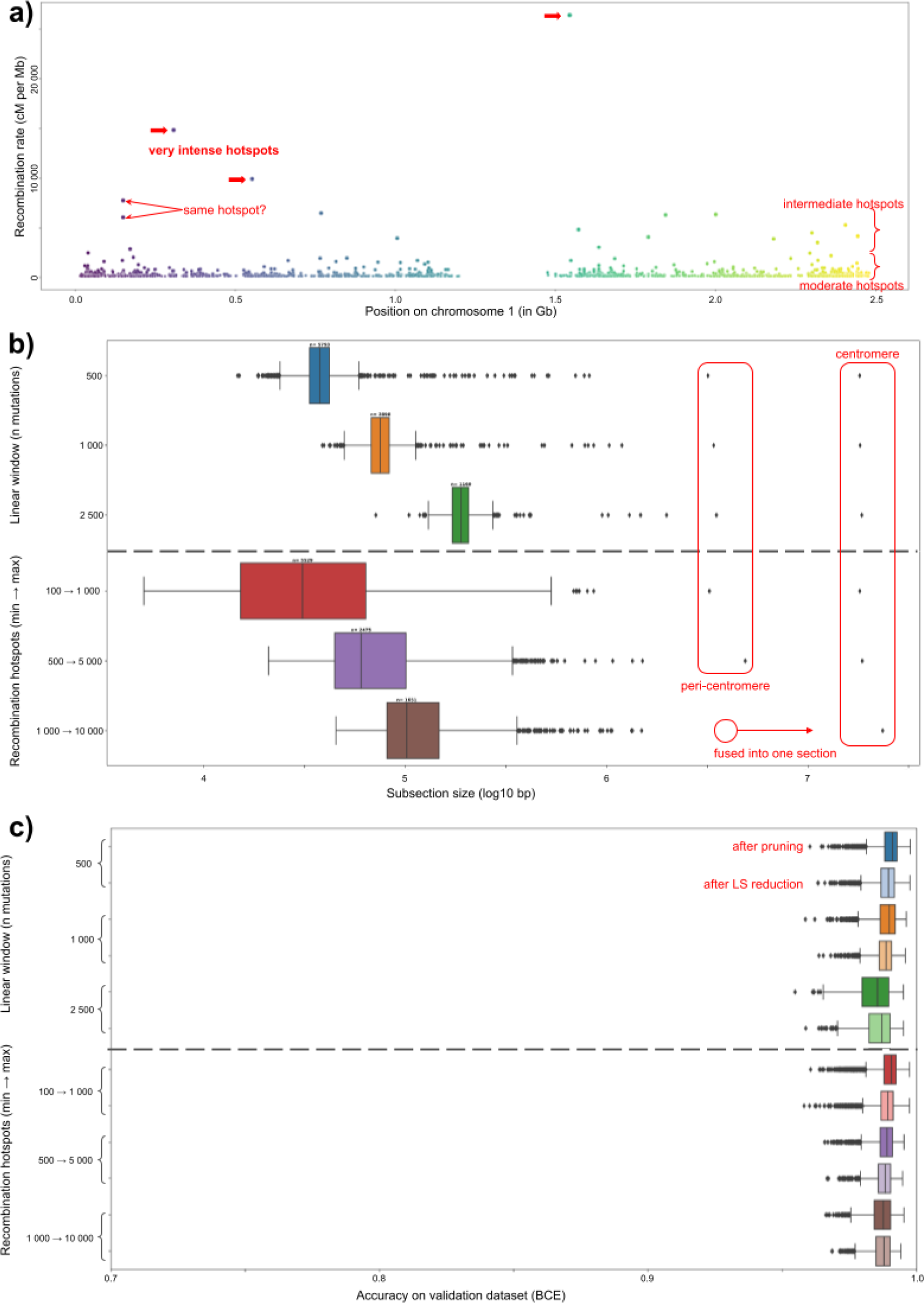
a)Scatter plot illustrating the distribution of recombination hotspots along Human chromosome 1. Vertical axis is the recombination rate at that hotspot. Some individual hotspots or categories of hotspots are highlighted manually. b)Boxplot of the size of genomic subsections depending on the segmentation method used. Interestingly, the centromere contains few listed mutations and therefore appears as one very large section. c)Boxplot showing the accuracy of autoencoder models depending on the segmentation method used (order is the same as figure 1a). For each pair of boxes, the richly coloured box is the original model with pruning implemented, and the lighter coloured box is a separate decoder trained based on a reduced latent space.

**Supplementary fig 2:**
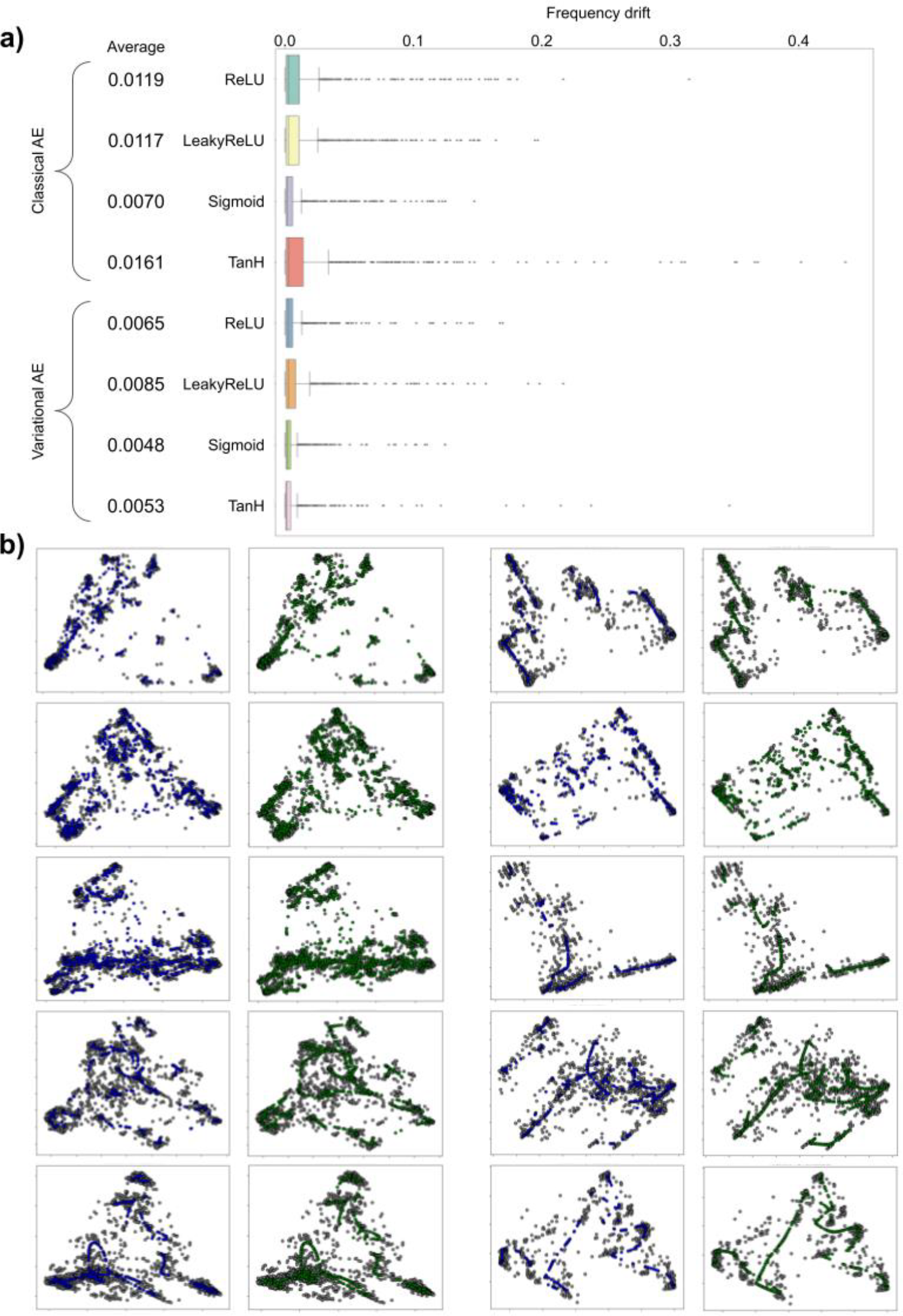
a)Boxplot of drift of each mutation for different autoencoder models. Average drift for each model is also included. b)Scatter plots of PCAs for different datasets as described for figure 2b, with each row illustrating two different sections. All genomic subsections studied in detail that were not shown in fig 2b are shown here.

**Supplementary fig 3:**
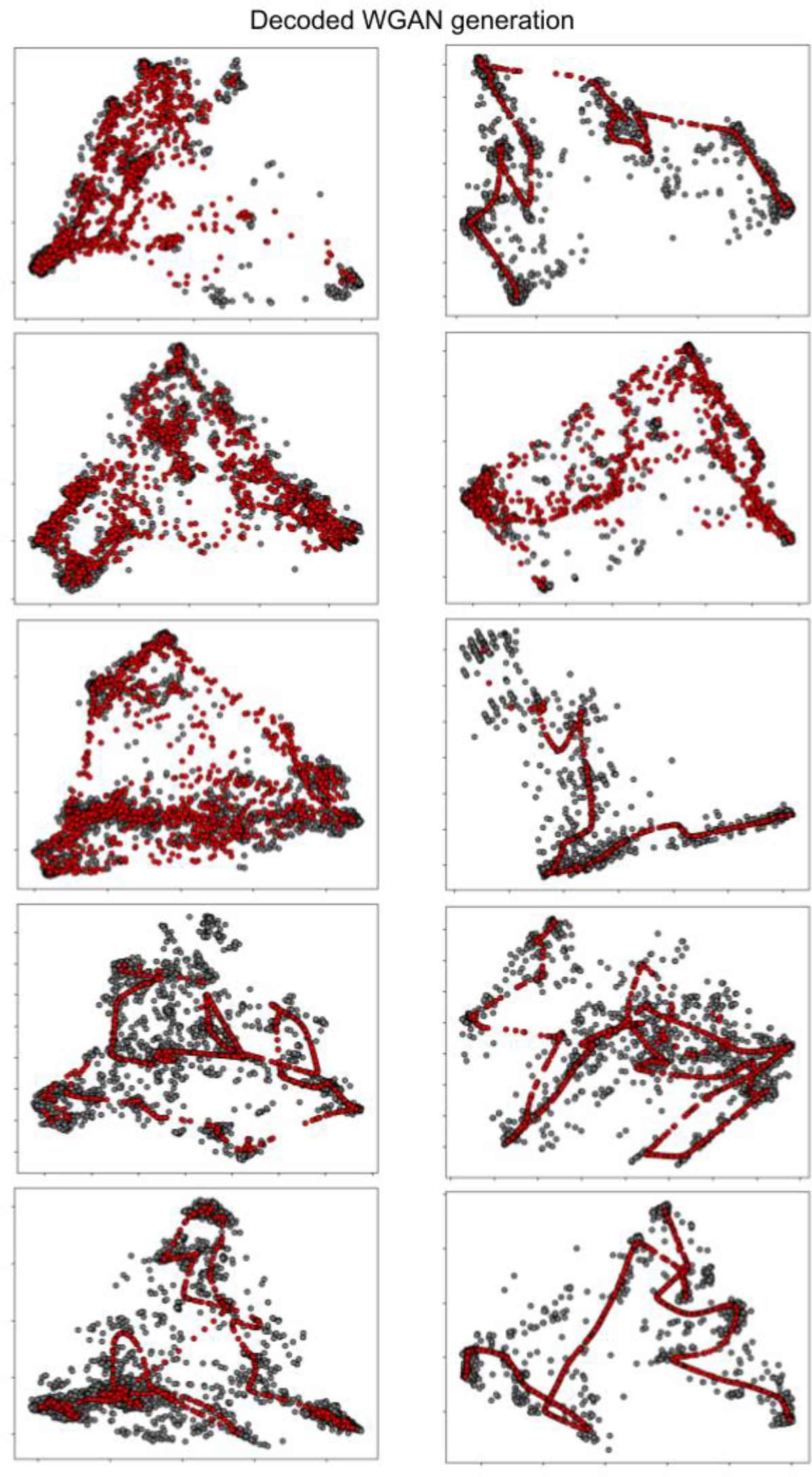
Scatter plots of PCAs for different datasets as described for figure 3a, with each row illustrating two different sections. All genomic subsections studied in detail that were not shown in fig 3a are shown here.

## Acknowledgments

C.B. wishes to thank the Ligue Contre le Cancer for providing financing for the doctoral thesis behind this project.

The authors thank Genotoul for providing data processing and storage resources for the machine learning models listed here.

## References

Arjovsky, Martin, Soumith Chintala, and Léon Bottou. 2017. ‘Wasserstein GAN’. arXiv. http://arxiv.org/abs/1701.07875.

Balloux, F. 2001. ‘EASYPOP (Version 1.7): A Computer Program for Population Genetics Simulations’. Journal of Heredity 92 (3): 301–2. 10.1093/jhered/92.3.301.

Carvajal-Rodríguez, Antonio. 2008. ‘GENOMEPOP: A Program to Simulate Genomes in Populations’. BMC Bioinformatics 9 (1): 223. 10.1186/1471-2105-9-223.

Halldorsson, Bjarni V., Gunnar Palsson, Olafur A. Stefansson, Hakon Jonsson, Marteinn T. Hardarson, Hannes P. Eggertsson, Bjarni Gunnarsson, et al. 2019. ‘Characterizing Mutagenic Effects of Recombination through a Sequence-Level Genetic Map’. Science 363 (6425): eaau1043. 10.1126/science.aau1043.

Hamming, R. W. 1950. ‘Error Detecting and Error Correcting Codes’. Bell System Technical Journal 29 (2): 147–60. 10.1002/j.1538-7305.1950.tb00463.x.

Joly, Yann, Ida Ngueng Feze, and Jacques Simard. 2013. ‘Genetic Discrimination and Life Insurance: A Systematic Review of the Evidence’. BMC Medicine 11 (January): 25. 10.1186/1741-7015-11-25.

Juan, Liran, Yongtian Wang, Jingyi Jiang, Qi Yang, Qinghua Jiang, and Yadong Wang. 2020. ‘PGsim: A Comprehensive and Highly Customizable Personal Genome Simulator’. Frontiers in Bioengineering and Biotechnology 8. https://www.frontiersin.org/articles/10.3389/fbioe.2020.00028.

Kim, Miran, and Kristin Lauter. 2015. ‘Private Genome Analysis through Homomorphic Encryption’. BMC Medical Informatics and Decision Making 15 (Suppl 5): S3. 10.1186/1472-6947-15-S5-S3.

Kingma, Diederik P., and Max Welling. 2013. ‘Auto-Encoding Variational Bayes’. arXiv. http://arxiv.org/abs/1312.6114.

Kramer, M.A. 1992. ‘Autoassociative Neural Networks’. Computers & Chemical Engineering 16 (4): 313–28. 10.1016/0098-1354(92)80051-A.

Kullback, S., and R. A. Leibler. 1951. ‘On Information and Sufficiency’. The Annals of Mathematical Statistics 22 (1): 79–86. 10.1214/aoms/1177729694.

Kuo, Tsung-Ting, and Anh Pham. 2021. ‘Detecting Model Misconducts in Decentralized Healthcare Federated Learning’. International Journal of Medical Informatics 158 (December): 104658. 10.1016/j.ijmedinf.2021.104658.

Lawson, Daniel John, Garrett Hellenthal, Simon Myers, and Daniel Falush. 2012. ‘Inference of Population Structure Using Dense Haplotype Data’. PLOS Genetics 8 (1): e1002453. 10.1371/journal.pgen.1002453.

Lowy-Gallego, Ernesto, Susan Fairley, Xiangqun Zheng-Bradley, Magali Ruffier, Laura Clarke, and Paul Flicek. 2019. ‘Variant Calling on the GRCh38 Assembly with the Data from Phase Three of the 1000 Genomes Project’. Wellcome Open Research 4 (December): 50. 10.12688/wellcomeopenres.15126.2.

Mu, John C., Marghoob Mohiyuddin, Jian Li, Narges Bani Asadi, Mark B. Gerstein, Alexej Abyzov, Wing H. Wong, and Hugo Y.K. Lam. 2015. ‘VarSim: A High-Fidelity Simulation and Validation Framework for High-Throughput Genome Sequencing with Cancer Applications’. Bioinformatics 31 (9): 1469–71. 10.1093/bioinformatics/btu828.

Peng, Bo, and Marek Kimmel. 2005. ‘simuPOP: A Forward-Time Population Genetics Simulation Environment’. Bioinformatics 21 (18): 3686–87. 10.1093/bioinformatics/bti584.

Price, W. Nicholson, and I. Glenn Cohen. 2019. ‘Privacy in the Age of Medical Big Data’. Nature Medicine 25 (1): 37–43. 10.1038/s41591-018-0272-7.

Prince, Anya E.R. 2017. ‘Insurance Risk Classification in an Era of Genomics: Is a Rational Discrimination Policy Rational?’ Nebraska Law Review 96 (3): 624–87.

Ramachandran, Prajit, Barret Zoph, and Quoc V. Le. 2017. ‘Searching for Activation Functions’. arXiv. http://arxiv.org/abs/1710.05941.

Schneider, Valerie A., Tina Graves-Lindsay, Kerstin Howe, Nathan Bouk, Hsiu-Chuan Chen, Paul A. Kitts, Terence D. Murphy, et al. 2017. ‘Evaluation of GRCh38 and de Novo Haploid Genome Assemblies Demonstrates the Enduring Quality of the Reference Assembly’. Genome Research 27 (5): 849–64. 10.1101/gr.213611.116.

Shabani, Mahsa, and Luca Marelli. 2019. ‘Re-Identifiability of Genomic Data and the GDPR’. EMBO Reports 20 (6): e48316. 10.15252/embr.201948316.

Tahmasbi, Rasool, and Matthew Keller. 2016. ‘GeneEvolve: A Fast and Memory Efficient Forward-Time Simulator of Realistic Whole-Genome Sequence and SNP Data’. Bioinformatics (Oxford, England) 33 (September). 10.1093/bioinformatics/btw606.

Tange, Ole. 2011. ‘GNU Parallel: The Command-Line Power Tool’. Login.

The 1000 Genomes Project Consortium. 2015. ‘A Global Reference for Human Genetic Variation’. Nature 526 (7571): 68–74. 10.1038/nature15393.

Yelmen, Burak, Aurélien Decelle, Linda Ongaro, Davide Marnetto, Corentin Tallec, Francesco Montinaro, Cyril Furtlehner, Luca Pagani, and Flora Jay. 2021. ‘Creating Artificial Human Genomes Using Generative Neural Networks’. PLOS Genetics 17 (2): e1009303. 10.1371/journal.pgen.1009303.

Yue, Jia-Xing, and Gianni Liti. 2019. ‘simuG: A General-Purpose Genome Simulator’. Bioinformatics 35 (21): 4442–44. 10.1093/bioinformatics/btz424.

